# Autologous antimicrobials affect transposon library composition

**DOI:** 10.64898/2026.07.07.736938

**Authors:** Biwen Wang, Zihao Teng, Tjalling Siersma, Frans van der Kloet, Leendert Hamoen

**Author notes:** These authors contributed equally (Biwen Wang, Zihao Teng).

## Abstract

Genome-wide transposon insertion sequencing (Tn-seq) is a powerful tool to measure the importance of genes for growth. In this study, we applied Tn-seq to the Gram-positive model system *Bacillus subtilis*, and found that after growth in liquid medium the transposon library lacked transposon insertions in several genes related to lipoteichoic acid biosynthesis and cell wall teichoic acid modification. This was unexpected since these genes are not essential for normal growth. By growing the transposon library as a confluent layer of cells, and as discrete colonies, we found that these genes are only important when the transposon library is grown as a confluent layer. Thus, growing the transposon library as a mixed population reduces the fitness of teichoic acid mutants. We provide further evidence showing that this anomaly is caused by the production of multiple autologous antimicrobials and/or toxins in the growth medium. These findings show that growing genome-wide mutant libraries as mixed cultures can influence the mutant library composition.

## INTRODUCTION

Transposon insertion sequencing (Tn-seq) is a powerful technique to determine the contribution of individual genes to cellular fitness on a genome-wide scale (Bourque et al., 2018; Cain et al., 2020; Kobras et al., 2021; Kreft et al., 2017). Key to the process is the construction of a saturated transposon library, ensuring multiple transposon insertions per gene, which is then grown under selective conditions, after which transposon insertion sites are identified by next-generation sequencing. The fitness of a gene is then calculated by comparing transposon insertion frequencies between the different conditions (van Opijnen et al., 2009; Willcocks et al., 2019; Zhang et al., 2022; Basta et al., 2025). For a complete genome-wide analysis, it is important that the transposon library fully covers all non-essential genes. While creating a transposon library with the Gram-positive model bacterium *Bacillus subtilis*, using the mariner transposon (Dobihal et al., 2022), we noticed that almost no transposon insertions were found in several lipoteichoic acid biosynthesis and teichoic acid modification genes. This was curious, since these genes are not important for growth. Here, we show that this anomaly is only observed when the library is grown as a mixed culture, and is caused by autologous antimicrobials accumulating in the medium. This finding is important for Tn-seq, CRISPRi-seq, and related techniques that employ genome-wide mutant libraries.

## RESULTS AND DISCUSSION

### Transposon library anomaly

To perform a Tn-seq analysis with *B. subtilis*, we employed an *in vitro* genomic transposon insertion protocol, using purified HimarC9 Mariner transposase (Johnson and Grossman, 2014; van Opijnen et al., 2009). The resulting transposon library was transformed into *B. subtilis* cells, yielding approximately 250,000 colonies that were pooled together and stored at −80 °C (Fig. 1A). To check the coverage of the transposon library, a frozen aliquot was diluted in LB medium to an OD_600_ of 0.2 and incubated at 37 °C with shaking for 1 h to allow recovery of cells from freezing, designated here as timepoint 1 (T1). Alternatively, the culture was first diluted to an OD_600_ of 0.006 and allowed to grow for ∼4 h until an OD_600_ of 1.6 was reached, corresponding to approximately 8 generations and designated as timepoint 2 (T2) (Fig. 1A). Chromosomal DNA from T1 and T2 was isolated for Tn-seq. The growth experiment was repeated and the Tn-seq data showed good reproducibility (Fig. S1).

**Fig. 1.**
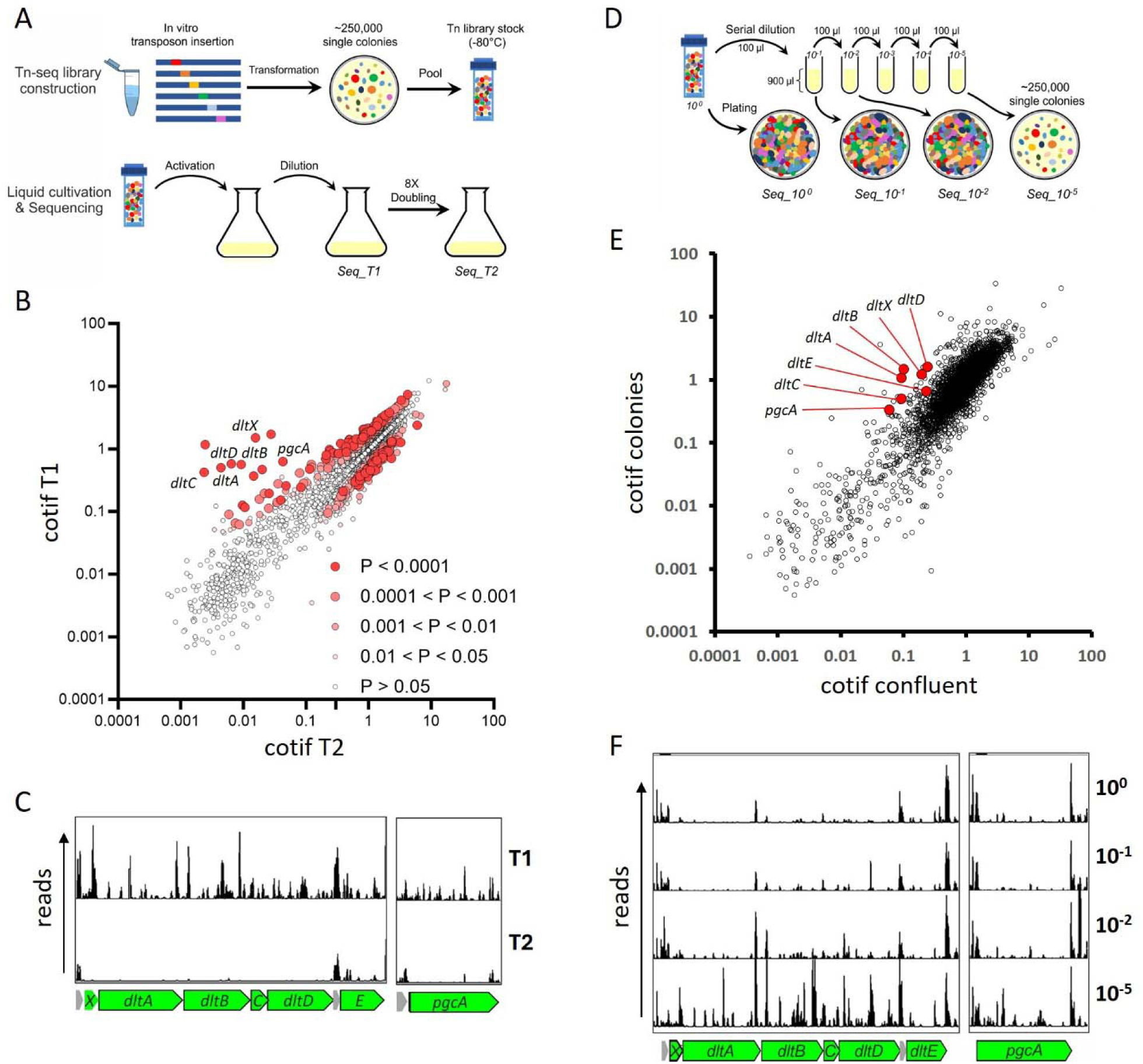
Comparison of transposon insertion frequencies of different library conditions. (A) *in vitro* transposon insertion and construction of the *B. subtilis* transposon library (upper panel), and subsequent dilution and growth to T1 and T2 (lower panel). (B) Scatter plot comparing cotif values obtained at T1 and T2. p-Values are colour coded based on fold-change calculations of cotif values. 142 essential genes had a cotif value of 0 and are not shown in the graph. (C) Transposon insertions mapped in *dltA* and *pgcA* loci for T1 and T2. (D) Schematic outline of the experiment whereby the transposon library is grown on solid nutrient agar at different dilutions. (E) Scatter plot comparing cotif values of the confluent growing plates with plates containing discrete colonies. Relevant genes mentioned in the main text are indicated. (F) Transposon hits mapped in *dltA* and *pgcA* loci from samples plated at different dilutions.

To determine gene fitness, the number of transposon insertions was divided by the theoretical number of insertions based on the number of TA sites in a gene (Table S1). For these <u>co</u>rrected transposon insertion frequency values, or cotif values, the first and last 10 % of each gene were discarded, since transposon insertions at either the very beginning or end are less likely to affect gene activity (van Opijnen et al., 2009). The advantage of cotif values is that a single timepoint can already provide useful information on gene fitness, and they help to interpret transposon insertion frequency differences between conditions in case of low insertion rates, since this can lead to unrealistically high fold-change differences. When the cotif values for both timepoints were plotted against each other in a scatter plot, we noticed a cluster of genes with normal cotif values for T1 but much lower values for T2 (Fig. 1B, C). The *dltXABCDE* operon is involved in D-alanylation of both cell wall teichoic acids and membrane-anchored lipoteichoic acids (Wecke et al., 1996), and *pgcA* is involved in lipoteichoic acid synthesis (Gründling and Schneewind, 2007; Matsuoka et al., 2016; Schirner et al., 2009). Other lipoteichoic acid synthesis genes, including *gtaB* and *ugtP*, showed significantly lower cotif values for T2, as well (Table S1). The lack of transposon insertions in these genes is surprising since none of them have been reported to be essential for growth, which we could confirm (Fig. S2).

### Effect of colony dilution on Tn-seq

A key difference between T1 and T2 was that in the former case cells were collected from transformation plates containing discrete colonies, whereas in the latter case, cells had grown for 8 generations in liquid culture. Possibly, the anomaly involving lipoteichoic acid synthesis and D-alanylation genes occurs when transposon mutants are grown as mixed populations. To test this, we made several dilutions of the Tn-library before plating on LB agar (Fig. 1D). With a 100,000-fold dilution, we obtained discrete colonies on the plates, whereas the undiluted, 10-fold, and 100-fold diluted samples grew as confluent layers. Cells were collected from the different plates for Tn-seq analysis (Table S2), and the cotif values from the confluent plates were compared with those of the 100,000-fold dilution plates (Fig. 1E). Clearly, mutants impaired in teichoic D-alanylation and lipoteichoic acid synthesis contain more transposon insertions when grown as discrete colonies (Fig. 1F).

### Competition in liquid medium

It has been reported that mutations in the *dlt* operon render *B. subtilis* more susceptible to the lipid II targeting lantibiotic nisin (Kingston et al., 2013). *B. subtilis* itself produces a number of antimicrobial compounds, including several bacteriocins, such as subtilosin, bacilysin and sublancin (Paik et al., 1998; Zheng et al., 1999). Possibly, the absence of either lipoteichoic acids or D-alanylation of teichoic acids makes cells vulnerable to their own secreted antimicrobials and toxins. To test this, we constructed a *dltA* deletion mutant by replacing the gene with an erythromycin resistance cassette (Koo et al., 2017), and grew this mutant to exponential phase. The culture was then mixed 1:1 with an exponentially growing wildtype culture, and the viable counts of both strains were determined after 1, 2, and 4 h by plating on nutrient agar with and without erythromycin. As shown in Fig. 2A, Δ*dltA* cells do not grow when mixed with the wildtype culture and start to die after 2 h. Comparable results were found for Δ*dltB*, Δ*dltC* and Δ*pgcA* (Fig. 2B).

**Fig. 2.**
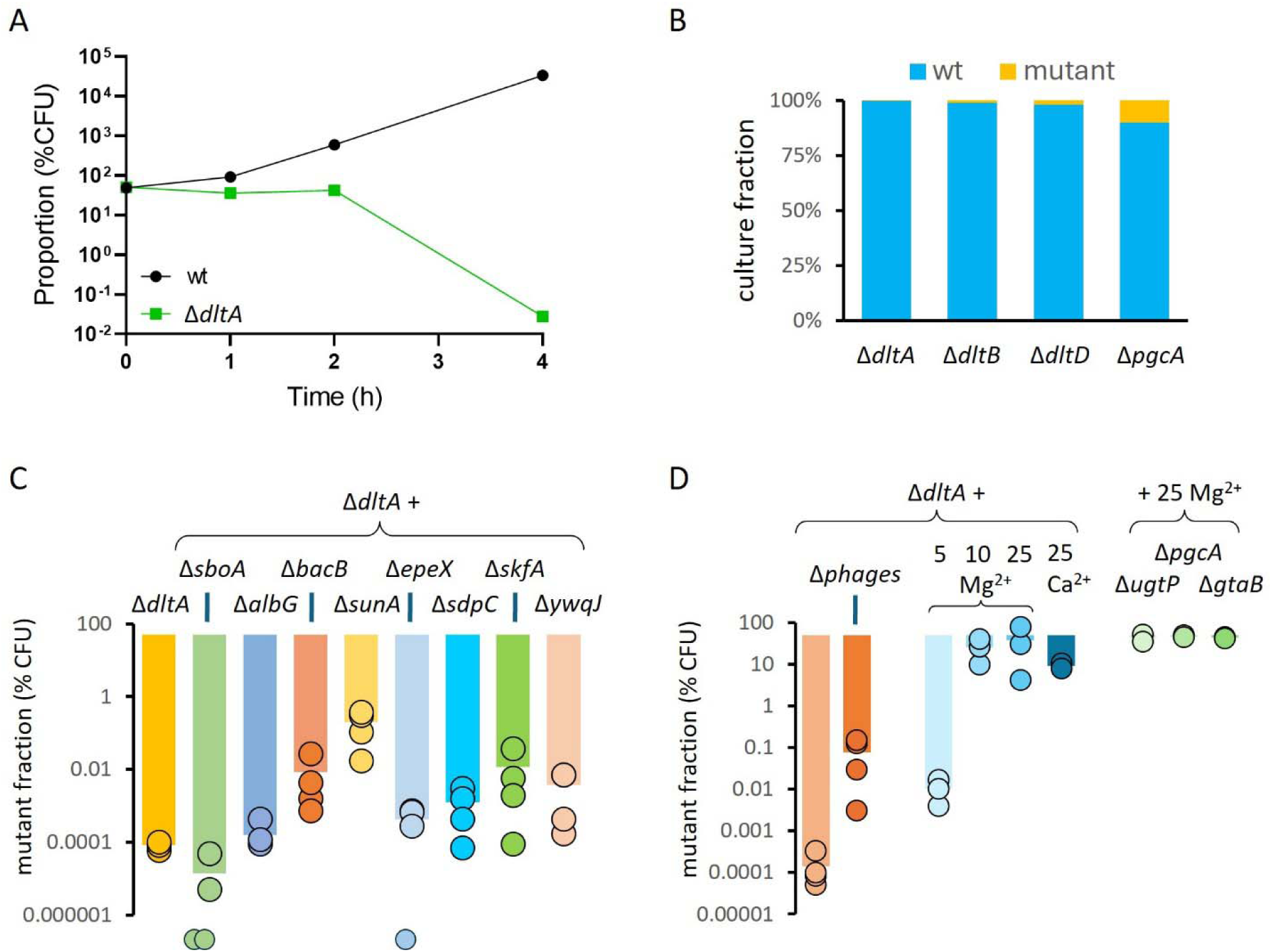
Competition disadvantage of teichoic acid mutants. (A) Growth competition between wildtype and Δ*dltA* mutant cells mixed and grown in LB medium. Surviving fractions were determined by colony forming units (CFU). (B) Fraction of different mutants after mixing at a 1:1 ratio with wildtype cells, and subsequent 4 h growth in LB medium. Start OD_600_ was approximately 0.006. (C) Survival of Δ*dltA* strains lacking different autologous antimicrobials or toxins after 4 h coculturing with wild type cells. See main text for explanation of mutants. (D) Left bars: Survival of a Δ*dltA* strain, and a Δ*dltA* lacking 6 prophages (Δphages), after 4 h coculturing with a Δphages strain with an intact *dltA* gene. Middle bars: Fraction of a Δ*dltA* strain after 4 h growth with the wildtype strain when either 5, 10, 25 mM Mg^2+^ or 25 mM Ca^2+^ is present in the LB medium. Right bars: Fraction of either the Δ*ugtP*, Δ*pgcA* or Δ*gtaB* strain after 4 h growth with a wildtype strain when 25 mM Mg^2+^ is present in the LB medium.

### Sensitivity of ΔdltA cells to autologous antimicrobials

These data strongly suggest that an intact teichoic acid cell wall layer protects *B. subtilis* cells against an autologous antimicrobial compound that accumulates in the culture, or possibly multiple autologous antimicrobials. *B. subtilis* produces a diversity of antimicrobials and toxins. The lab strain 168 lacks the cofactor to synthesize non-ribosomal peptides and polyketides (Mootz et al., 2001; Nakano et al., 1992; Tsuge et al., 1999). However, this strain still produces the bacteriocin subtilosin, the dipeptide antibiotic bacilysin, and the glycocin sublancin (Paik et al., 1998; Zheng et al., 1999). To examine whether these antibiotics are responsible for the competitive disadvantage of the Δ*dltA* mutant, we introduced mutations that block the synthesis of subtilosin (Δ*sboA*, Δ*albG*), bacilysin (Δ*bacB*) or sublancin (Δ*sunA*) (Paik et al., 1998; Steinborn et al., 2005; Zheng et al., 2000). We then mixed each antibiotic mutant with the corresponding strain that also carried a Δ*dltA* mutation, and measured the viable counts after 4 h of incubation. As shown in Fig. 2C, preventing the expression of subtilosin (Δ*sboA*, Δ*albG*) did not increase the fitness of a Δ*dltA* mutant. On the other hand, preventing the expression of bacilysin (Δ*bacB*) or sublancin (Δ*sunA*) resulted in incremental fitness improvements (Fig. 2C). However, because the fractions are shown on a logarithmic scale, removal of either still resulted in a > 99 % reduction of Δ*dltA* cells in the mixed cultures.

*B. subtilis* produces several toxic peptides, including EpeX, SdpC and SkfA (Popp et al., 2020; González-Pastor et al., 2003; Koskiniemi et al., 2013). Deleting *epeX* had a marginal effect on the sensitivity of the Δ*dltA* mutant, and although deleting either *skfA* or *sdpC* had a stronger effect, again the overall result was that most Δ*dltA* cells were killed after 4 h of competition (Fig. 2C). *B. subtilis* also possesses several toxin-antitoxin systems, some of which are secreted, including YwqJ (Kobayashi, 2021). Blocking expression of this toxin also had a minimal effect on the fitness of Δ*dltA* cells (Fig. 2C). These findings suggest that the reduced fitness is caused by multiple autologous antimicrobial compounds. Consistent with this idea, a genome-reduced *B. subtilis* strain, lacking all known antimicrobial peptides and toxins (Reuß et al., 2017), showed no fitness difference when cocultured with an otherwise identical strain carrying a *dltA* mutation (data not shown).

### Contribution of prophages

As a final control, we examined the role of bacteriophages. Teichoic acids function as receptor molecules for many bacteriophages (Räisänen et al., 2007; Addo et al., 2024). *W*e used a *B. subtilis* strain cured of all six prophage elements (Reuß et al., 2016), and introduced the *dltA* deletion into this strain. When the resulting strain was mixed with the original Δ6 prophage strain, its viable count was still very low after 4 h of competition (Fig. 2D).

### Effect of divalent cations

The addition of D-alanines to the polyglycerolphosphate chain of teichoic acids reduces their negative charge, leading to an increase in the density of the cell wall (Saar-Dover et al., 2012). In fact, the increased sensitivity of a Δ*dltA* mutant to the antibiotic methicillin can be suppressed by the addition of magnesium ions (Wecke et al., 1997). Because of this, and the fact that LB is low in divalent cations (Wee and Wilkinson, 1988; Pallares-Vega et al., 2021), we tested whether the addition of extra Mg^2+^ or Ca^2+^ ions could influence the competition between wildtype and Δ*dltA* cells. To this end, the coculturing experiments were repeated but now in the presence of 5, 10, or 25 mM MgCl_2_, or 25 mM CaCl_2_. Interestingly, the addition of 10-25 mM of these ions almost completely restored the fitness of the Δ*dltA*, Δ*ugtP* and Δ*gtaB* mutants (Fig. 2D).

## Conclusion

Here we have shown that antimicrobials and/or toxins secreted by *B. subtilis* can affect the composition of a transposon library, because cells require proper teichoic acid cell wall polymers to protect themselves against these compounds. We were unable to pinpoint this effect to a particular antimicrobial compound or toxin, and we assume that the reduced fitness is caused by multiple autologous antimicrobial compounds. Since the vast majority of bacteria produce antimicrobials and toxins, the effect of autologous secreted antimicrobials and toxins should be considered when aiming to construct a comprehensive mutant library for Tn-seq, CRISPRi-seq, and related studies.

## MATERIAL AND METHODS

### Bacterial strains, cultivation conditions and mutant construction

Bacterial strains used in this study are listed in Table S3. The tryptophan-prototrophic wildtype *B. subtilis* strain BSB1 was used as the parent wildtype strain (Nicolas et al., 2012). Cells were grown in Luria-Bertani medium (LB), containing 10 g/L Tryptone, 5 g/L Yeast Extract, and 10 g/L NaCl. When necessary, plates or media were supplemented with 150 µg/ml spectinomycin, 5Sµg/ml erythromycin (*erm*) or 5Sµg/ml chloramphenicol (cm) for selection.

For *B. subtilis* transformation, cells were grown in Spizizen minimal medium (SMM), containing 15SmM (NH_4_)_2_SO_4_, 80SmM K_2_HPO_4_, 44SmM KH_2_PO_4_, 3SmM tri-sodium citrate, 0.5 % glucose, 6SmM MgSO, 0.2SmgSml^−1^ tryptophan, 0.02 % casamino acids, and 0.00011 % ferric ammonium citrate ((NH_4_)_5_Fe(C_6_H_4_O_7_)_2_) (Spizizen and J, 1958). Natural competence was induced following the procedure described in (Anagnostopoulos and Spizizen, 1961). Single knockout mutants were constructed by transferring chromosomal DNA from the genomic mutant BKE library (Koo et al., 2017) into the BSB1 background following the standard transformation procedure as described in (Anagnostopoulos and Spizizen, 1961).

### Preparation of transposon library

Construction of the *B. subtilis* transposon insertion library and transposon insertion sequencing (Tn-seq) were carried out essentially as previously described (Johnson and Grossman, 2014; van Opijnen et al., 2014). For a single 20 μl *in vitro* transposition reaction, 6 μg genomic DNA of BSB1, 2 μg transposon-harboring plasmid pCJ4, 0.4 μg in-house purified mariner transposase Himar1-C9, and 10 μl 2x buffer A (41 mM HEPES pH 7.9, 19 % glycerol, 187 mM NaCl, 19 mM MgCl_2_, 476 μg/ml BSA, 3.8 mM dithiothreitol) were combined and incubated overnight at 30 °C. The reaction product was precipitated by adding 4 μl 3 M sodium acetate, pH 5.2 and 100 μl ice-cold ethanol and centrifuged for 30 min at 16000 RCF and 4 °C. The DNA pellet was washed twice with ice-cold 70 % ethanol, air-dried and resuspended in the aqueous system containing 2 μl 10x buffer B (500 mM tris-Cl pH 7.8, 100 mM MgCl_2_, 10 mM dithiothreitol), 2 μl 1 mg/ml BSA and 11 μl water. Next, 4 μl 2.5 mM dNTPs and 0.5 μl T4 DNA Polymerase (NEB) were added, and the mixture was incubated for 20 min at 12 °C to repair transposon insertion junctions. After heat inactivation of the polymerase at 75 °C for 15 min, 0.2 μl 2.6 mM NAD and 0.5 μl *E. coli* DNA ligase (NEB) were added and incubated at 16 °C overnight to repair the nicked DNA strands before storage at −20 °C. BSB1 cells were grown in Spizizen’s minimal medium (SMM) to induce genetic competence as described previously (Anagnostopoulos and Spizizen, 1961; Hamoen et al., 2002). Upon reaching maximum competence, cells were 6-fold concentrated, 5 μl of Tn-gDNA was added to 250 μl concentrated competent cells and incubated at 37°C with vigorous shaking. After 60 min of incubation, 250 μl LB was added and incubation continued for 30 min, and cells were plated on LB agar plates supplemented with 150 μg/ml spectinomycin and grown at 37 °C overnight. Using Tn-gDNA from twelve 20 μl reactions, we obtained approximately 250,000 individual transformants per transformation. Colonies were scraped from the plate using SMM medium and pooled together (30 ml, OD_600_ of 125). After homogenization by thorough vortexing, cells were gently spun down at 3000 RCF for 8 min, and the pellet was resuspended in 20 ml SMM containing 15 % glycerol. The library was aliquoted into 1 ml portions, flash frozen in liquid nitrogen, and stored at −80 °C.

### Growth conditions for Tn-seq

For the first Tn-seq experiment, a 1 ml aliquot of the transposon library was quickly thawed and diluted to an OD_600_ of 0.2 using prewarmed LB. Cells were allowed to recover for 1 h at 37°C with vigorous shaking (210 rpm). Then, the culture was further diluted in 100 ml prewarmed LB to an OD_600_ of 0.00625 (T1), and the culture was grown for approximately eight doubling periods, reaching an OD_600_ of 1.6 (T2). Cells were sampled for genomic DNA extraction.

For the second Tn-seq experiment, an aliquot of the transposon library was thawed, and 200 μl was plated on LB agar supplemented with spectinomycin (10 cm x 10 cm plate). The remainder of the library was subjected to a 10-fold serial dilution in LB medium before plating. The undiluted, 10^-1^ and 10^-2^ diluted samples grew as confluent layers, whereas the 10^-5^ dilution formed discrete colonies on the plates. Around 250,000 single colonies from the 10^-5^ dilution plates were scraped and pooled together for genomic DNA extraction. Cells from the undiluted, 10^-1^, and 10^-2^ dilution plates were also scraped separately for the same subsequent procedures.

### Tn-seq sequencing library construction

The purified genomic DNA from the Tn library was subjected to MmeI digestion by adding 3 μg gDNA, 3 μl MmeI (NEB), 0.44 μl 32 mM S-adenosylmethionine and 20 μl CutSmart buffer in a 200 μl total reaction volume, followed by incubation for 2.5 h at 37 °C. Subsequently, 2 μl calf intestinal phosphatase (NEB) was added for 1 h at 37 °C. DNA was cleaned up with phenol/chloroform/isoamyl alcohol mixture (PCI, 25:24:1), precipitated by sodium acetate and ethanol, and washed twice with 70 % ice-cold ethanol. DNA pellets were resuspended in 27 μl 2 mM tris-Cl pH 8.5, and 2 μl was used for DNA concentration measurement. Meanwhile, a customized Illumina sequencing R1 adapter was annealed using complementary oligonucleotides BW-TnR1-S and BW-TnR1-AntiS, 200 μM in 1 mM Tris-HCl pH 8.5, by heating 95 °C for 5 min, followed by cooling to 4 °C at a ramp rate of −1 °C/min. 1 μl R1 adapter was added to 25 μl DNA, together with 1 μl T4 DNA ligase (NEB) and 3 μl 10× T4 DNA ligase buffer, followed by an overnight incubation at 16 °C. The adapter-ligated DNA junctions were purified using 21 μl (0.7x volume AMPure XP beads (Beckman Coulter)), according to the manufacturer’s protocol. Briefly, DNA was eluted from beads with 33 μl MQ water and 2 μl of the elution was used to determine the yield using a Qubit fluorometer (Thermo Fisher). Purified adapter-ligated DNA was used as a template for PCR amplification of Illumina sequencing libraries using a universal primer BW-TnPCR-Uni and a specific barcode primer (e.g., BW-Tn-BC30) and Q5® Hot Start High-Fidelity 2x Master Mix (NEB). PCR cycling conditions were as follows. Step-1: 98 °C for 2 min. Step-2: 98 °C for 10 sec. Step-3: 68 °C for 25 sec. Step-4: 72 °C for 10 sec. Repeat Steps 2-4 for 14-18 cycles. Step-5: 72 °C for 2 min. Then hold at 4 °C. PCR products were separated on an 8 % non-urea PAGE gel, and DNA bands of approximately 165 bp were excised. Gel slices were crushed and incubated with 400 μl DNA extraction buffer (300 mM NaCl, 1.0 mM EDTA, 10 mM Tris-HCl pH 8.0) in 2 ml tubes in a rolling rack at 4 °C overnight. Polyacrylamide was removed through a SpinX column, and DNA was precipitated with 1 ml ice cold ethanol and 1 μl GlycoBlue coprecipitant (Thermo Fisher) for at least 2 h at −80 °C. Precipitated material was collected by 30 min centrifugation at 20,000 RCF and 4 °C. Pellets were washed twice with ice cold 80 % ethanol, and resuspended in 0.1x TE buffer (NEB). The DNA concentration was measured using a Qubit fluorometer. Tn-seq libraries were sequenced using an Illumina NextSeq 550 System with NextSeq 500/550 High Output v2.5 kit (75-bp read length). All primers are listed in Table S4.

### Tn-seq data analysis

The raw sequence data were processed using the web-based platform Galaxy in combination with a customized Python package called *map_functions*. Briefly, *Trimmomatic* was used to trim Illumina adapter sequences and filter bad reads. *Cutadapt* was used to remove the transposon inverted repeat (IR) sequence ACAGGTTGGATGATAAGT from the 3’-end. Reads that did not contain the IR sequence were discarded and only reads of 12-20 bp size were kept. Reads were aligned to a *B. subtilis* reference genome (GenBank accession no.NC_000964.3) using *Bowtie2*, resulting in a set of alignments in SAM format. We used the in-house Python package *map_functions* to extract mapping information from the SAM files based on TA dinucleotide locations. A TA definition CSV file was created from the genome FASTA and annotation GFF3 files, identifying 218,025 unique TA sites across the genome. Mapped forward reads (direction = 0) that ended with TA, and reverse reads (direction = 16) that started with TA were kept, and reads that did not start or end with TA were removed from the SAM file. Intergenic regions make up only 10 % of the genome, but are AT rich and contain 18 % of potential HimarC9 insertion sites. However, insertions into these regions are less likely to affect gene fitness. In addition, it has been shown that transposon insertions close to the beginning or the end of a gene are less likely to affect gene activity (Jacobs et al., 2003; van Opijnen et al., 2009). Therefore, only reads that mapped into known genetic features, based on BSU locus_tags, were counted. Insertions located within 10 % (open reading frame length) of the 5’ or 3’ end of each genetic feature were removed. In total, there are 183,963 TA sites located in the 4420 loci of the *B. subtilis* genome. After removal of TA sites within 10 % of either the 5’- or 3’-end, 142,642 TA sites in 4409 loci were left. Transposon insertion numbers for the Tn-seq experiments are listed in the table of Fig. S3.

To assign fitness values to the different loci, we determined the <u>co</u>rrected transposon insertion frequency (cotif) values by normalizing the read numbers, and calculating the theoretical read numbers per gene based on total reads and the number of TA sites in a gene. The cotif values were then obtained by dividing the actual reads by the theoretical reads.

### Growth rate measurement

Growth rates were measured in shake flask cultivation. Briefly, overnight cultures of wildtype and mutants were prepared in 10 ml LB medium in 100 ml flasks and incubated at 30 °C with 210 rpm shaking. 30 °C was used, instead of 37 °C, to prevent sporulation. 5 μg/ml erythromycin was added for selection of mutants. The next morning, the overnight cultures were diluted to an OD_600_ of 0.2 in 10 ml fresh LB medium and grown at 37 °C for 1 h. Then the cultures were again diluted in 10 ml LB medium, now to an OD_600_ of 0.00625, and the OD_600_ was followed over time.

### Growth competition assay

Growth competition assays were performed as follows. Overnight cultures of wildtype and mutants were prepared in 10 ml LB medium in 100 ml flasks and incubated at 30 °C with 210 rpm shaking. 5 μg/ml erythromycin was added for mutant culturing. The next morning, the cultures were diluted to an OD_600_ of 0.2 in 10 ml fresh LB medium and grown at 37 °C for 1 h. The OD_600_ was then measured, and the two competing cultures were mixed 1:1, and the mixture was diluted in 10 ml LB medium to an OD_600_ of 0.00625. The mixed culture was grown for 4 h at 37 °C and 210 rpm, resulting in approximately 8 generations. After 4 h, samples were serially diluted and plated on LB agar containing erythromycin or chloramphenicol. The colony forming units (CFU) on plates with and without erythromycin/chloramphenicol were used to determine the numbers of wild-type and mutant cells, respectively.

## ACKNOWLEDGMENT

We would like to thank Gaurav Dugar (DNA&RNA interaction lab, University of Amsterdam) for his advice and inspiring discussions, Samantha Zoomer and Mara Hoogteijling for initial experiments with mutant competition, Berit Kooter for initial experiments with minimal genome mutants, Rob Dekker and Selina van Leeuwen (MAD, University of Amsterdam) for providing excellent sequencing services, and statistical support. This research was partially funded by CSC China Scholarship Council fellowships.

## Author contributions

Biwen Wang: Conceptualization, Data curation, Formal analysis, Investigation, Validation, Visualization, Writing original draft & editing. Zihao Teng: Conceptualization, Data curation, Formal analysis, Investigation, Validation, Writing & editing. Tjalling Siersma: Formal analysis, Investigation. Frans van der Kloet: Data curation, Formal analysis, Software. Leendert Hamoen: Conceptualization, Formal analysis, Funding acquisition, Project administration, Resources, Writing & editing.

## Data availability

All data supporting the findings of this study are presented in the paper, Supplementary Information or Source Data file. Strains are available upon request. The Tn-seq raw reads and processed tabular files have been submitted to the Gene Expression Omnibus (GEO) with accession number GSE322624.

## REFERENCE

Addo MA, Zang Z, Gerdt JP. 2024. Chemical inhibition of cell surface modification sensitizes bacteria to phage infection. RSC Chemical Biology 5:1132–1139. DOI: 10.1039/D4CB00070F

Anagnostopoulos C, Spizizen J. 1961. Requirements for transformation in *Bacillus subtilis*. Journal of bacteriology 81:741.

Basta DW, Campbell IW, Sullivan EJ, Hotinger JA, Hullahalli K, Garg M, Waldor MK. 2025. Inducible transposon mutagenesis identifies bacterial fitness determinants during infection in mice. Nature Microbiology 10:1171–1183. DOI: 10.1038/s41564-025-01975-z

Bourque G, Burns KH, Gehring M, Gorbunova V, Seluanov A, Hammell M, Imbeault M, Izsvák Z, Levin HL, Macfarlan TS, Mager DL, Feschotte C. 2018. Ten things you should know about transposable elements. Genome Biology 19:199. DOI: 10.1186/s13059-018-1577-z

Cain AK, Barquist L, Goodman AL, Paulsen IT, Parkhill J, van Opijnen T. 2020. A decade of advances in transposon-insertion sequencing. Nature Reviews Genetics 21:526–540. DOI: 10.1038/s41576-020-0244-x

Dobihal GS, Flores-Kim J, Roney IJ, Wang X, Rudner DZ. 2022. The WalR-WalK Signaling Pathway Modulates the Activities of both CwlO and LytE through Control of the Peptidoglycan Deacetylase PdaC in *Bacillus subtilis*. Journal of Bacteriology 204:e00533–21. DOI: 10.1128/jb.00533-21

González-Pastor JE, Hobbs EC, Losick R. 2003. Cannibalism by sporulating bacteria. Science 301:510–513. DOI: 10.1126/science.1086462, PMID: 12817086

Gründling A, Schneewind O. 2007. Genes required for glycolipid synthesis and lipoteichoic acid anchoring in *Staphylococcus aureus*. Journal of bacteriology 189:2521–2530. DOI: 10.1128/JB.01683-06, PMID: 17209021

Hamoen LW, Smits WK, Jong A de, Holsappel S, Kuipers OP. 2002. Improving the predictive value of the competence transcription factor (ComK) binding site in *Bacillus subtilis* using a genomic approach. Nucleic Acids Research 30:5517–5528. DOI: 10.1093/nar/gkf698

Jacobs MA, Alwood A, Thaipisuttikul I, Spencer D, Haugen E, Ernst S, Will O, Kaul R, Raymond C, Levy R, Chun-Rong L, Guenthner D, Bovee D, Olson MV, Manoil C. 2003. Comprehensive transposon mutant library of *Pseudomonas aeruginosa*. Proceedings of the National Academy of Sciences 100:14339–14344. DOI: 10.1073/pnas.2036282100

Johnson CM, Grossman AD. 2014. Identification of host genes that affect acquisition of an integrative and conjugative element in *Bacillus subtilis*. Molecular Microbiology 93:1284–1301. DOI: 10.1111/MMI.12736, PMID: 25069588

Kingston AW, Liao X, Helmann JD. 2013. Contributions of the σW, σM and σX regulons to the lantibiotic resistome of *Bacillus subtilis*. Molecular Microbiology 90:502–518. DOI: 10.1111/mmi.12380

Kobayashi K. 2021. Diverse LXG toxin and antitoxin systems specifically mediate intraspecies competition in *Bacillus subtilis* biofilms. PLOS Genetics 17:e1009682. DOI: 10.1371/journal.pgen.1009682

Kobras CM, Fenton AK, Sheppard SK. 2021. Next-generation microbiology: from comparative genomics to gene function. Genome Biology 22:123. DOI: 10.1186/s13059-021-02344-9

Koo B-M, Kritikos G, Farelli JD, Todor H, Tong K, Kimsey H, Wapinski I, Galardini M, Cabal A, Peters JM. 2017. Construction and analysis of two genome-scale deletion libraries for *Bacillus subtilis*. Cell systems 4:291–305.

Koskiniemi S, Lamoureux JG, Nikolakakis KC, t’Kint de Roodenbeke C, Kaplan MD, Low DA, Hayes CS. 2013. Rhs proteins from diverse bacteria mediate intercellular competition. Proceedings of the National Academy of Sciences 110:7032–7037. DOI: 10.1073/pnas.1300627110

Kreft J-U, Plugge CM, Prats C, Leveau JHJ, Zhang W, Hellweger FL. 2017. From Genes to Ecosystems in Microbiology: Modeling Approaches and the Importance of Individuality. Frontiers in Microbiology 8. DOI: 10.3389/fmicb.2017.02299

Matsuoka S, Seki T, Matsumoto K, Hara H. 2016. Suppression of abnormal morphology and extracytoplasmic function sigma activity in *Bacillus subtilis ugtP* mutant cells by expression of heterologous glucolipid synthases from *Acholeplasma laidlawii*. Bioscience, Biotechnology, and Biochemistry 80:2325–2333. DOI: 10.1080/09168451.2016.1217147

Mootz HD, Finking R, Marahiel MA. 2001. 4’-phosphopantetheine transfer in primary and secondary metabolism of *Bacillus subtilis*. The Journal of biological chemistry 276:37289–37298. DOI: 10.1074/JBC.M103556200, PMID: 11489886

Nakano MM, Corbell N, Besson J, Zuber P. 1992. Isolation and characterization of sfp: a gene that functions in the production of the lipopeptide biosurfactant, surfactin, in *Bacillus subtilis*. Molecular and General Genetics MGG 232:313–321. DOI: 10.1007/BF00280011

Nicolas P, Mäder U, Dervyn E, Rochat T, Leduc A, Pigeonneau N, Bidnenko E, Marchadier E, Hoebeke M, Aymerich S, Becher D, Bisicchia P, Botella E, Delumeau O, Doherty G, Denham EL, Fogg MJ, Fromion V, Goelzer A, Hansen A, Härtig E, Harwood CR, Homuth G, Jarmer H, Jules M, Klipp E, Le Chat L, Lecointe F, Lewis P, Liebermeister W, March A, Mars RAT, Nannapaneni P, Noone D, Pohl S, Rinn B, Rügheimer F, Sappa PK, Samson F, Schaffer M, Schwikowski B, Steil L, Stülke J, Wiegert T, Devine KM, Wilkinson AJ, Maarten van Dijl J, Hecker M, Völker U, Bessières P, Noirot P. 2012. Condition-Dependent Transcriptome Reveals High-Level Regulatory Architecture in *Bacillus subtilis*. Science 335:1103–1106. DOI: 10.1126/science.1206848

Paik SH, Chakicherla A, Hansen JN. 1998. Identification and Characterization of the Structural and Transporter Genes for, and the Chemical and Biological Properties of, Sublancin 168, a Novel Lantibiotic Produced by Bacillus subtilis 168. Journal of Biological Chemistry 273:23134–23142. DOI: 10.1074/jbc.273.36.23134, PMID: 9722542

Pallares-Vega R, Macedo G, Brouwer MSM, Hernandez Leal L, van der Maas P, van Loosdrecht MCM, Weissbrodt DG, Heederik D, Mevius D, Schmitt H. 2021. Temperature and Nutrient Limitations Decrease Transfer of Conjugative IncP-1 Plasmid pKJK5 to Wild *Escherichia coli* Strains. Frontiers in Microbiology 12. DOI: 10.3389/fmicb.2021.656250

Popp PF, Benjdia A, Strahl H, Berteau O, Mascher T. 2020. The Epipeptide YydF Intrinsically Triggers the Cell Envelope Stress Response of *Bacillus subtilis* and Causes Severe Membrane Perturbations. Frontiers in Microbiology 11. DOI: 10.3389/fmicb.2020.00151

Räisänen L, Draing C, Pfitzenmaier M, Schubert K, Jaakonsaari T, von Aulock S, Hartung T, Alatossava T. 2007. Molecular Interaction between Lipoteichoic Acids and *Lactobacillus delbrueckii* Phages Depends on d-Alanyl and α-Glucose Substitution of Poly(Glycerophosphate) Backbones. Journal of Bacteriology 189:4135–4140. DOI: 10.1128/jb.00078-07

Reuß DR, Altenbuchner J, Mäder U, Rath H, Ischebeck T, Sappa PK, Thürmer A, Guérin C, Nicolas P, Steil L, Zhu B, Feussner I, Klumpp S, Daniel R, Commichau FM, Völker U, Stülke J. 2017. Large-scale reduction of the *Bacillus subtilis* genome: Consequences for the transcriptional network, resource allocation, and metabolism. Genome Research 27:289–299. DOI: 10.1101/GR.215293.116/-/DC1, PMID: 27965289

Reuß DR, Thürmer A, Daniel R, Quax WJ, Stülke J. 2016. Complete Genome Sequence of *Bacillus subtilis subsp. subtilis* Strain Δ6. Genome Announcements 4:10.1128/genomea.00759-16. DOI: 10.1128/genomea.00759-16

Saar-Dover R, Bitler A, Nezer R, Shmuel-Galia L, Firon A, Shimoni E, Trieu-Cuot P, Shai Y. 2012. D-Alanylation of Lipoteichoic Acids Confers Resistance to Cationic Peptides in Group B *Streptococcus* by Increasing the Cell Wall Density. PLOS Pathogens 8:e1002891. DOI: 10.1371/journal.ppat.1002891

Schirner K, Marles-Wright J, Lewis RJ, Errington J. 2009. Distinct and essential morphogenic functions for wall- and lipo-teichoic acids in *Bacillus subtilis*. The EMBO Journal 28:830. DOI: 10.1038/EMBOJ.2009.25, PMID: 19229300

Spizizen J, J S. 1958. Transformation of Biochemically Deficient Strains of *Bacillus subtilis* by deoxyribonucleate. Proceedings of the National Academy of Sciences 44:1072–1078. DOI: 10.1073/pnas.44.10.1072, PMID: 16590310

Steinborn G, Hajirezaei M-R, Hofemeister J. 2005. *bac* genes for recombinant bacilysin and anticapsin production in *Bacillus* host strains. Archives of Microbiology 183:71–79. DOI: 10.1007/s00203-004-0743-8

Tsuge K, Ano T, Hirai M, Nakamura Y, Shoda M. 1999. The genes degQ, pps, and lpa-8 (sfp) are responsible for conversion of *Bacillus subtilis* 168 to plipastatin production. Antimicrobial agents and chemotherapy 43:2183–2192. DOI: 10.1128/AAC.43.9.2183, PMID: 10471562

van Opijnen T, Bodi KL, Camilli A. 2009. Tn-seq: high-throughput parallel sequencing for fitness and genetic interaction studies in microorganisms. Nature Methods 6:767–772. DOI: 10.1038/nmeth.1377

van Opijnen T, Lazinski DW, Camilli A. 2014. Genome-wide fitness and genetic interactions determined by Tn-seq, a high-throughput massively parallel sequencing method for microorganisms. Current Protocols in Molecular Biology 106:7.16.1-7.16.24. DOI: 10.1002/0471142727.mb0716s106

Wecke J, Madela K, Fischer W. 1997. The absence of D-alanine from lipoteichoic acid and wall teichoic acid alters surface charge, enhances autolysis and increases susceptibility to methicillin in *Bacillus subtilis*. Microbiology 143:2953–2960. DOI: 10.1099/00221287-143-9-2953

Wecke J, Perego M, Fischer W. 1996. D-alanine deprivation of *Bacillus subtilis* teichoic acids is without effect on cell growth and morphology but affects the autolytic activity. Microbial Drug Resistance 2:123–129. DOI: 10.1089/mdr.1996.2.123, PMID: 9158734

Wee S, Wilkinson BJ. 1988. Increased outer membrane ornithine-containing lipid and lysozyme penetrability of *Paracoccus denitrificans* grown in a complex medium deficient in divalent cations. Journal of Bacteriology 170:3283–3286. DOI: 10.1128/jb.170.7.3283-3286.1988

Willcocks S, Huse KK, Stabler R, Oyston PCF, Scott A, Atkins HS, Wren BW. 2019. Genome-wide assessment of antimicrobial tolerance in *Yersinia pseudotuberculosis* under ciprofloxacin stress. Microbial Genomics 5:e000304. DOI: 10.1099/mgen.0.000304

Zhang Y, Li Z, Xu X, Peng X. 2022. Transposon mutagenesis in oral *Streptococcus*. Journal of Oral Microbiology 14:2104951. DOI: 10.1080/20002297.2022.2104951

Zheng G, Hehn R, Zuber P. 2000. Mutational Analysis of the *sbo-alb* Locus of *Bacillus subtilis*: Identification of Genes Required for Subtilosin Production and Immunity. Journal of Bacteriology 182:3266–3273. DOI: 10.1128/jb.182.11.3266-3273.2000

Zheng G, Yan LZ, Vederas JC, Zuber P. 1999. Genes of the *sbo-alb* Locus of *Bacillus subtilis* Are Required for Production of the Antilisterial Bacteriocin Subtilosin. Journal of Bacteriology 181:7346–7355. DOI: 10.1128/jb.181.23.7346-7355.1999

