## Supplementary Information for "Autologous antimicrobials affect transposon library composition"

**Content:**

- **Legend Table S1 (Excel file):** Tn-seq data and cotif values for samples T1 and T2
- **Legend Table S2 (Excel file):** Tn-seq data and cotif values for plates with different dilutions
- **Table S3:** Strains used in this study
- **Table S4:** Primers used in this study
- **Fig. S1:** Scatter plots of Tn-insertion reads per gene for the biological replicates
- **Fig. S2:** Growth rates of different teichoic acid mutants
- **Fig. S3:** Transposon insertion data from the different Tn-seq experiments
- **References**

**Legend Table S1 (Excel file). Tn-seq data and cotif values for samples T1 and T2**

| Description of columns in Table S1 | |
| --- | --- |
| locus_tag | unique identifier |
| geneLength | length of gene in nucleotides |
| nr_TA_sites | number of TA sites in a gene |
| TA_position | TA dinucleotide positions |
| geneLength_5'3'-10% | length of gene in nucleotides minus 10% of both sides |
| nr_TA_5'3'-10% | number of TA sites in a gene minus 10% of both sides |
| measured read_T1i | number of transposon insertions per gene time point T1, replicate i |
| measured read_T1ii | number of transposon insertions per gene time point T1, replicate ii |
| measured read_T2i | number  of transposon insertions per gene time point T2, replicate i |
| measured read_T2ii | number of transposon insertions per gene time point T2, replicate ii |
| cotif_T1 | cotif value at time point T1 based on the two replicates |
| cotif_T2 | cotif value at time point T2 based on the two replicates |
| cotif_T2/T1 | fold-change cotif value calculated by dividing cotif value T2 by cotif value T1 |
| pVal_T2/T1 | related p-value |
| gene | gene name |
| function | gene function |
| description | gene description |
| essential | indication whether a gene is essential or not |

**Legend Table S2 (Excel file). Tn-seq data and cotif values for plates with different dilutions.**

| Description of columns in Table S2 | |
| --- | --- |
| locus_tag | unique identifier |
| geneLength | length of gene in nucleotides |
| nr_TA_sites | number of TA sites in a gene |
| TA_position | TA dinucleotide positions |
| geneLength_5'3'-10% | length of gene in nucleotides minus 10% of both sides |
| nr_TA_5'3'-10% | number of TA sites in a gene minus 10% of both sides |
| reads_10^0 | number of transposon insertions per gene for the undiluted library |
| reads_10^-1 | number of transposon insertions per gene for the 10x diluted library |
| reads_10^-2 | number of transposon insertions per gene for the 100x diluted library |
| reads_10^-5 | number of transposon insertions per gene for the 100,000x diluted library |
| cotif_10^0 | cotif value for the undiluted library |
| cotif_10^-1 | cotif value for the 10x diluted library |
| cotif_10^-2 | cotif value for the 100x diluted library |
| cotif_(10^0&-1&-2) | average cotif values of undiluted, 10x and 100x diluted libraries (all confluent) |
| cotif_10^-5 | cotif value for the 100,000x diluted library |
| gene | gene name |
| function | gene function |
| description | gene description |
| essential | indication whether a gene is essential or not |

**Table S3. Strains used in this study.**

| **Strain** | **Genotype** | **Source or reference** |
| --- | --- | --- |
| BSB1 | *B. subtilis wildtype 168 trp+* | Lab strain |
| *BS1* | *Δphages, Δ6 (∆SPß, ∆skin, ∆ PBSX, ∆ proф1, ∆ pks, ∆ proф3) trp+* | Lab strain |
| BKE38500 | *dltA::erm*, *trpC2* | (Koo et al., 2017) |
| BKE38510 | *dltB::erm*, *trpC2* | (Koo et al., 2017) |
| BKE38520 | *dltC::erm*, *trpC2* | (Koo et al., 2017) |
| BKE38530 | *dltD::erm*, *trpC2* | (Koo et al., 2017) |
| BKE09310 | *pgcA::erm*, *trpC2* | (Koo et al., 2017) |
| BKE21920 | *ugtP::erm, trpC2* | (Koo et al., 2017) |
| BKE35670 | *gtaB::erm, trpC2* | (Koo et al., 2017) |
| BKE37350 | *sboA::erm, trpC2* | (Koo et al., 2017) |
| BKE37430 | *albG::erm, trpC2* | (Koo et al., 2017) |
| BKE37730 | *bacB::erm, trpC2* | (Koo et al., 2017) |
| BKE21480 | *sunA::erm, trpC2* | (Koo et al., 2017) |
| BKE40180 | *epeX::erm, trpC2* | (Koo et al., 2017) |
| BKE33770 | *sdpC::erm, trpC2* | (Koo et al., 2017) |
| BKE01910 | *skfA::erm, trpC2* | (Koo et al., 2017) |
| BKE36190 | *ywqJ::erm* | (Koo et al., 2017) |
| BWSZ03 | *BSB1, dltA::erm* | This study |
| BWSZ04 | *BSB1, dltB::erm* | This study |
| BWSZ05 | *BSB1, dltC::erm* | This study |
| BWSZ06 | *BSB1, dltD::erm* | This study |
| BWSZ08 | *BSB1, pgcA::erm* | This study |
| BWB168 | *BSB1, ugtP::ery* | This study |
| BWB169 | *BSB1, gtaB::ery* | This study |
| BWB170 | *BS1, dltA::ery* | This study |
| BWB189 | *BSB1, dltA::cm,* *epeX::erm* | This study |
| BWB191 | *BSB1, dltA::cm,* *ywqJ::erm* | This study |
| BWB292 | *BSB1, dltA::cm, sboA::erm* | This study |
| BWB293 | *BSB1, dltA::cm,* albG::erm | This study |
| BWB294 | *BSB1, dltA::cm,* *bacB::erm* | This study |
| BWB295 | *BSB1, dltA::cm,* *sunA::erm* | This study |
| BWB297 | *BSB1, dltA::cm,* *sdpC::erm* | This study |
| BWB307 | *BSB1, dltA::cm,* *skfA::erm* | This study |

**Table S4. Primers used in this study.**

| **name** | | **Sequence 5’-3’** |
| --- | --- | --- |
| TnR1-S | TCTTTCCCTACACGACGCTCTTCCGATCTNN | |
| TnR1-AntiS | /5Phos/AGATCGGAAGAGCGTCGTGTAGGGAAAGA/3Phos/ | |
| Tn-Uni | AATGATACGGCGACCACCGAGATCTACACTCTTTCCCTACACGACGCTCTTCCGATCT | |
| Tn-BC30 | CAAGCAGAAGACGGCATACGAGATCCGGTGGTGACTGGAGTTCAGACGTGTGCTCTTCCGATCTAGACCGGGGACTTATCATCCAACCTGT | |
| Tn-BC37 | CAAGCAGAAGACGGCATACGAGATATTCCGGTGACTGGAGTTCAGACGTGTGCTCTTCCGATCTAGACCGGGGACTTATCATCCAACCTGT | |
| Tn-BC38 | CAAGCAGAAGACGGCATACGAGATAGCTAGGTGACTGGAGTTCAGACGTGTGCTCTTCCGATCAGACCGGGGACTTATCATCCAACCTGT | |
| Tn-BC39 | CAAGCAGAAGACGGCATACGAGATGTATAGGTGACTGGAGTTCAGACGTGTGCTCTTCCGATCAGACCGGGGACTTATCATCCAACCTGT | |
| Tn-BC40 | CAAGCAGAAGACGGCATACGAGATTGGATCACGTGACTGGAGTTCAGACGTGTGCTCTTCCGATCAGACCGGGGACTTATCATCCAACCTGT | |
| Tn-BC41 | CAAGCAGAAGACGGCATACGAGATGTCGTCGTGACTGGAGTTCAGACGTGTGCTCTTCCGATCTAGACCGGGGACTTATCATCCAACCTGT | |
| Tn-BC43 | CAAGCAGAAGACGGCATACGAGATGCTGTAGTGACTGGAGTTCAGACGTGTGCTCTTCCGATCAGACCGGGGACTTATCATCCAACCTGT | |
| Tn-BC44 | CAAGCAGAAGACGGCATACGAGATATTATAGTGACTGGAGTTCAGACGTGTGCTCTTCCGATCAGACCGGGGACTTATCATCCAACCTGT | |
| Tn-BC45 | CAAGCAGAAGACGGCATACGAGATGAATGAGTGACTGGAGTTCAGACGTGTGCTCTTCCGATCAGACCGGGGACTTATCATCCAACCTGT | |
| Tn-BC46 | CAAGCAGAAGACGGCATACGAGATTCGGGAGTGACTGGAGTTCAGACGTGTGCTCTTCCGATCAGACCGGGGACTTATCATCCAACCTGT | |
| Tn-BC47 | CAAGCAGAAGACGGCATACGAGATCTTCGAGTGACTGGAGTTCAGACGTGTGCTCTTCCGATCAGACCGGGGACTTATCATCCAACCTGT | |
| Tn-BC48 | CAAGCAGAAGACGGCATACGAGATTGCCGAGTGACTGGAGTTCAGACGTGTGCTCTTCCGATCAGACCGGGGACTTATCATCCAACCTGT | |

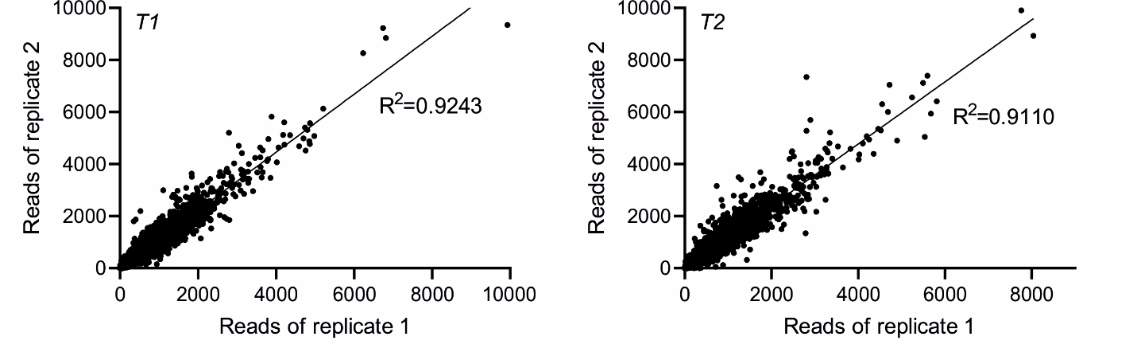
**Fig. S1**

**Fig. S1. Scatter plots of Tn-insertion reads per gene for the biological replicates.**

Scatter plots comparing the transposon insertion reads per gene for the two replicates at each time point (T1 and T2). Correlations are expressed as coefficients of determination (R^2^).

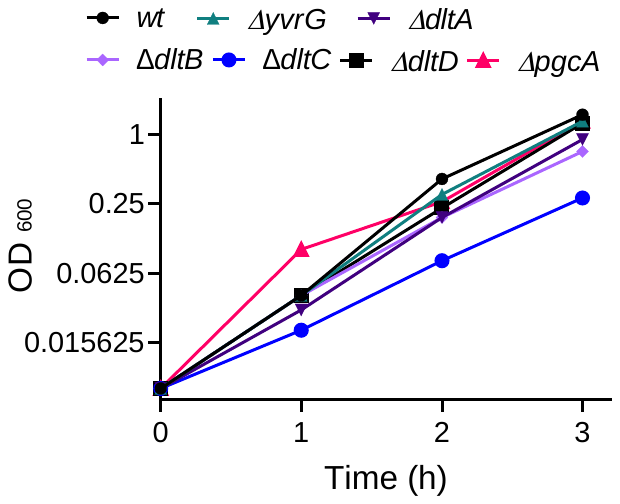
**Fig. S2**

**Fig. S2. Growth rates of different teichoic acid mutants.**

Growth was measured in LB medium by monitoring the optical density at 600 nm. The different mutants are described in the main text.

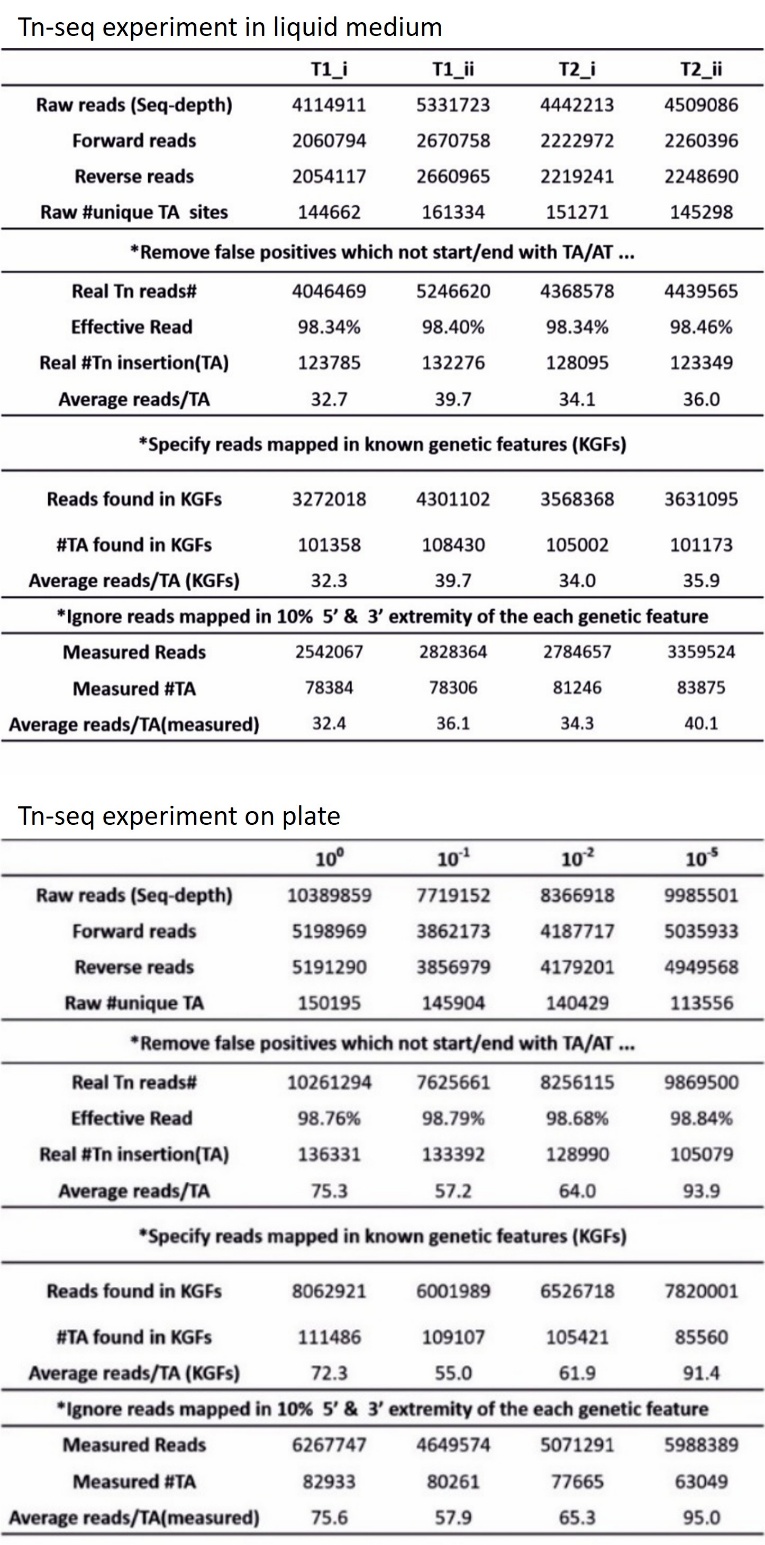
**Fig. S3**

**Fig. S3. Transposon insertion data from the different Tn-seq experiments.**

Summary of TA and transposon read numbers after subsequent processing steps.

**References**

Koo B-M, Kritikos G, Farelli JD, Todor H, Tong K, Kimsey H, Wapinski I, Galardini M, Cabal A, Peters JM. 2017. Construction and analysis of two genome-scale deletion libraries for *Bacillus subtilis*. Cell systems **4**:291–305.
